# Isolation of functional endogenous clathrin-coated vesicles and in vitro reconstitution of pure clathrin cages

**DOI:** 10.64898/2026.11.03.763552

**Authors:** Yinchao Wang, Xianshengjie Lang, Keqin Wu, Jinyue Li, Chuning Yao, Wenfei Li, Shuangli Sun

## Abstract

Clathrin-coated vesicles (CCVs) mediate key endocytic trafficking events, and dysfunction of CCVs is linked to multiple human diseases. Conventional centrifugation-based purification protocols can enrich endogenous CCVs but fail to eliminate intraluminal smooth vesicles (SVs) and co-isolated exosomes; these lipid contaminants reduce sample homogeneity and hinder high-resolution structural and proteomic analysis. Here, we present an integrated workflow to isolate biologically active endogenous CCVs from porcine brain and reconstitute homogeneous, SV-free clathrin cages. Crude CCVs are first enriched by differential and sucrose density gradient centrifugation to deplete the majority of exosomes. Clathrin cages are then depolymerized under alkaline conditions, allowing separation of soluble clathrin and adaptor proteins from SVs and residual lipid contaminants via ultracentrifugation. Purified coat proteins are reassembled in vitro into uniform clathrin cages. Functional assays confirm that purified CCVs retain Auxilin-Hsc70-dependent uncoating activity, and the enclosed SVs carry active vATPase proton pumps. This approach produces well-defined samples suitable for cryo-electron microscopy and mass spectrometry-based investigation of clathrin assembly and disassembly.

## 1. INTRODUCTION

Clathrin-coated vesicles (CCVs) are protein-enveloped vesicles in cells, closely associated with various human diseases. CCV formation is initiated by clathrin, which binds to the plasma membrane to form clathrin-coated pits (CCPs). Then clathrin triskelia and adaptor proteins (APs) are sequentially recruited to bend the membrane. Upon CCPs maturation, the GTPase dynamin is recruited and assembles into short helical rings around the necks of the CCPs to catalyze membrane fission, leading to the release of fully formed CCVs. The lumen of newly generated CCVs encloses smooth vesicles (SVs, synaptic vesicles) as cargo. After fulfilling trafficking functions, CCVs undergo uncoating mediated by Auxilin and Hsc70, which disassembles the outer clathrin cage and releases soluble clathrin and APs, while the internal SVs retain their integrity. The Vacuolar-type ATPases (vATPase) on SVs generates a membrane potential, energizing neurotransmitter release[1-5]. Ultimately, SVs fuse with the target membrane structure to mediate the transmembrane delivery of cargo molecules and orchestrate intercellular signaling. (Figure 1). Dysregulation of clathrin-dependent trafficking is linked to multiple neurological disorders including Alzheimer’s disease, Parkinson’s disease and epilepsy, as well as cancer and immune dysfunction [6–8].

**Figure 1.**
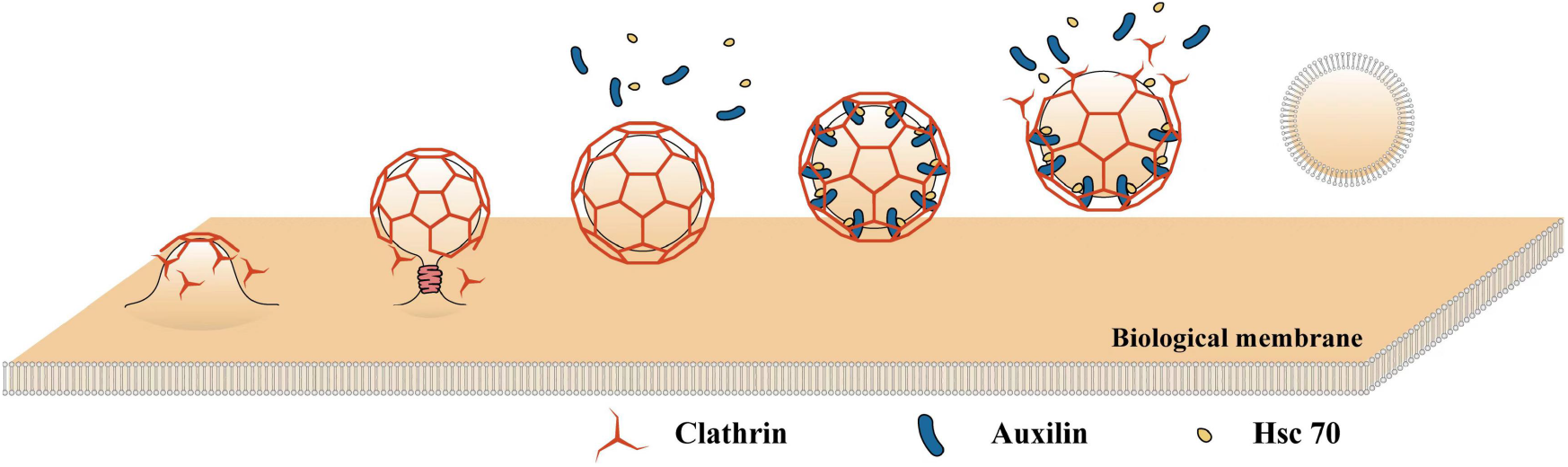
Membrane trafficking and uncoating cycle of clathrin-coated vesicles (CCVs). Schematic illustration showing the sequential assembly of clathrin-coated pits, membrane scission by dynamin to generate CCVs, and subsequent uncoating driven by Auxilin and Hsc70, which releases the internal smooth vesicle (SV). Clathrin, Auxilin and Hsc70 are indicated by red, blue and yellow symbols, respectively.

To dissect CCV biology and uncover its roles in disease-associated membrane trafficking pathways, reliable purification of endogenous CCVs with minimal contamination is a prerequisite. Multiple purification workflows have been developed. The prevailing strategy combines differential centrifugation with density gradient centrifugation to enrich endogenous CCVs from mammalian tissues [9, 10]. However, these methods rely on the sedimentation coefficient of CCVs, which results in the co-sedimentation of exosomes and other extracellular vesicles due to their similar size and density to CCVs.This extrinsic vesicular contamination creates a major confounding factor: it is difficult to clearly distinguish whether the observed biochemical or functional signals specifically to CCVs rather than copurified exosomes.

The demand for highly purified endogenous CCVs continues to grow, driven by both structural biology and proteomic investigations. In proteomic studies, sample purity directly determines the reliability of protein identification. Adaptor proteins (APs) are essential, low-abundance components governing CCV biogenesis [11]. Early work only detected relatively abundant AP variants such as AP180; the adoption of Ficoll-sucrose gradient centrifugation improved CCV yields and permitted the identification of additional trace APs including AP1, AP2 and AP50 [12,13]. For structural biology, impurities reduces sample homogeneity and degrade data quality, limiting the final resolution. Native endogenous CCVs are intrinsically heterogeneous due to fluid lipid bilayers and uneven coat protein distribution. Critically, internal SVs encapsulated within CCVs further interfere with cryo-EM reconstruction. To resolve the atomic-level mechanisms of clathrin coat assembly and disassembly, these SV contaminants must be eliminated, and clathrin subunits need to be reassembled in vitro into uniform clathrin cages. Therefore, depolymerization followed by controlled reconstitution represents an indispensable strategy to generate homogeneous clathrin cages suitable for reliable proteomic identification and high-resolution cryo-EM analysis.

Here we describe a novel workflow to obtain high-purity endogenous CCVs and reconstruct clathrin cages in vitro. We first enrich crude CCVs by differential and density gradient centrifugation. The isolated CCVs are then depolymerized to release clathrin and adaptor proteins, which enables the removal of exosomes and intraluminal SV contaminants. The purified coat proteins are reassembled in vitro to form uniform clathrin-coated cages. In vitro experiments demonstrate that the endogenous CCVs prepared by this workflow preserve their physiological function and enzymatic activity. This approach yields high-quality samples that support high-precision studies using structural biology and mass spectrometry-based proteomics.

## 2. Materials and Methods

### 2.1 Buffers

1. 10X buffer A: 250 mM HEPES, 1.25 M Potassium acetate, 50 mM Magnesium acetate, pH=7.0 (Note 1)
2. 1X buffer A: 25 mM Hepes, 125 mM Potassium acetate, 5 mM Magnesium acetate, pH=7.0.
3. Buffer B: To prepare 6.25% (m/V) Ficoll-sucrose solution, 1.25 g Ficoll and 1.25 g sucrose are dissolved in 1X buffer A to 20 ml (Note 2).
4. Buffer C: 2 M Tris, 2 mM EDTA, unadjusted pH.
5. Buffer D: 20 mM triethanolamine, 1 mM EDTA, pH = 8.0.
6. Buffer E: 100 mM MES, 15 mM Magnesium chloride, 0.2 mM EGTA, pH=6.5.
7. 20% (m/V) sucrose solution: Consists of 20% (m/V) sucrose and 1X buffer A (Note 2).
8. 40% (m/V) sucrose solution: Consists of 20% (m/V) sucrose and 1X buffer A (Note 2).
9. 20%-40% (m/V) continuous sucrose density gradient solution (Note 2):
  a. Add 7 ml 20% sucrose solution to the Beckman 14*89mm round bottom thin wall opening ultracentrifuge tube.
  b. Add 7 ml 40% sucrose solution to the bottom of the 20% sucrose solution Slowly. An apparent boundary between the two solutions could be seen.
  c. Cover with a 1 ml pulg. Prepare continuous sucrose density gradient solution (20%-40% m/V) with a BioComp Model 153 Gradient Station IP.

All solutions are stored at 4 °C and on ice during extraction.

### 2.2 Preparation of homogenization

Obtain eight fresh pig brains, each weighing approximately 95 g, stored in an ice box (Figure 2A). Using forceps, carefully remove as much fascia as possible from the surface of the pig brains (Note 4). Next, homogenize the pig brains using 1X Buffer A (approximately 50 ml per 100g of tissue) in a homogenizer until the mixture reaches a uniform, milkshake-like consistency (Note 5) (Figure 2B).

**Figure 2.**
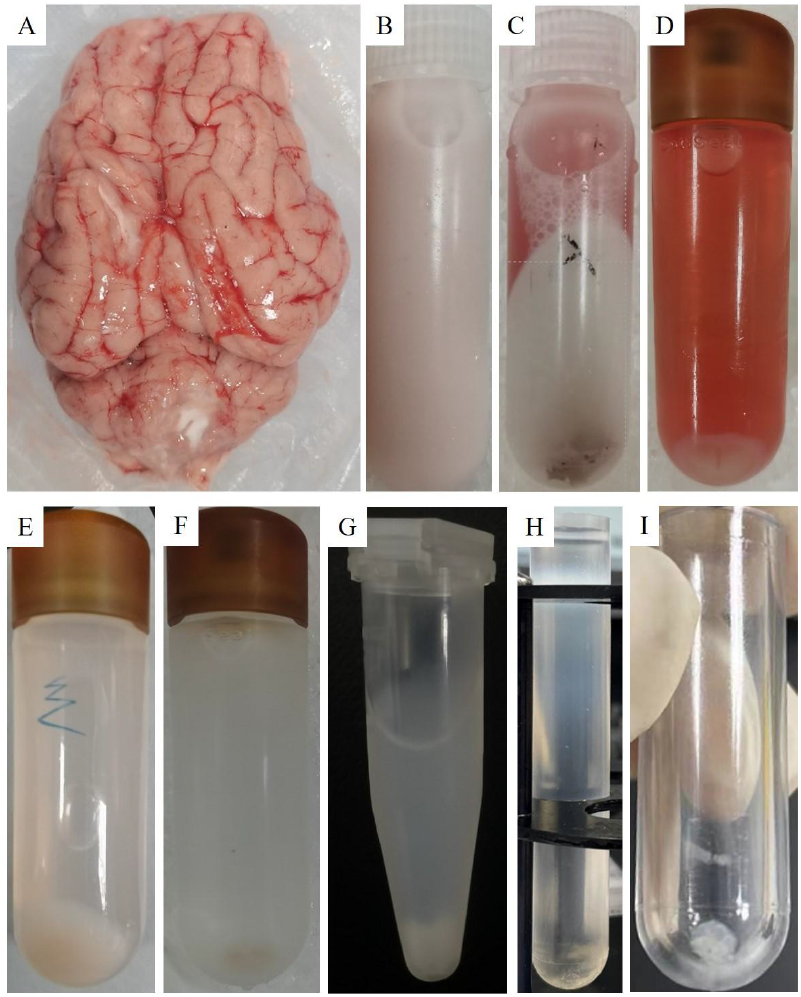
Overview of the CCV extraction and clathrin cage reconstruction workflow. Photographs showing key intermediate samples obtained throughout the purification procedure. (A) Fresh porcine brain tissue. (B) Brain tissue homogenate. (C-H) Sample appearance after successive centrifugation steps during CCV enrichment. (I) Final pellet of in vitro reconstructed clathrin cages.

### 2.3 Precipitation of large impurities with poor solubility

Aliquot the homogenized sample into centrifuge tubes and centrifuge at 12,000 g for 40 minutes at 4°C. It is important to note that the pellet will be loose and may tend to float (Figure 2C), so the supernatant (S1) should be carefully aspirated (Note 6). The pellet is then resuspended in Buffer A to the original volume and collected as sample P1.

### 2.4 Precipitation of lipid fractions

Transfer the supernatant into six Beckman 26 x 77 mm OptiSeal tubes and centrifuge at 56,000 g for 60 minutes at 4°C using a Beckman Optima-Max-XP ultracentrifuge with an MLA-50 rotor. Collect the supernatant as sample S2 (Figure 2D). Discard the remaining supernatant. Resuspend the pellet thoroughly with 10 ml of 1X Buffer A by gently tapping with a 1 ml pipette. Then, add an equal volume of Buffer B to the resuspended pellet and mix thoroughly. Pass the mixture through a syringe fitted with a 25-gauge needle, applying gentle and even pressure. The resulting mixture is collected as sample P2.

### 2.5 Enrichment of vesicles

Aliquot the mixture into a Beckman 26 x 77 mm OptiSeal tube and centrifuge at 43,000 g for 20 minutes at 4°C using a Beckman Optima-Max-XP ultracentrifuge with an MLA-50 rotor (Note 7). Collect the supernatant as sample S3 (Figure 2E). Resuspend the pellet in an equal volume of Buffer A and collect it as sample P3. To the supernatant, add 1X Buffer A to dilute it to approximately 180 ml and mix thoroughly. Aliquot the diluted supernatant into six Beckman 26 x 77 mm OptiSeal tubes and ultracentrifuge at 91,000 g for 60 minutes at 4°C using a Beckman Optima-Max-XP ultracentrifuge with an MLA-50 rotor. Collect the supernatant as sample S4 (Figure 2F). Resuspend the pellet in 2 ml of Buffer A and pass it through a syringe with a 25-gauge needle, applying gentle and even pressure. This is collected as sample P4 (Note 8).

### 2.6 Removal of the cytoskeleton

Aliquot the resuspended sample into two 1.5 ml EP tubes and centrifuge at 18,700 g for 10 minutes at 4°C. Collect the supernatant as sample S5 (Figure 2G). Resuspend the pellet in an equal volume of Buffer A and collect it as sample P5.

### 2.7 Isolation of exosomes

Prepare 20% and 40% sucrose solutions as described above. Carefully layer the supernatant on top of the continuous sucrose density gradient. A clear boundary between the sample and the sucrose solution should be visible. Centrifuge at 145,000 g for 90 minutes at 4°Cusing a Beckman Optima XPN-100 ultracentrifuge with an SW-41 rotor (Figure 2H). Separate the samples (0.5 ml per fraction) using a GILSON FC 203B fraction collector. The UV absorption curve should show two peaks at 20%-22.5% and 22.5%-24.5% sucrose densities. Collect the second peak, which contains the CCVs (Note 9). Fractions containing CCVs are stained with uranyl acetate for Transmission Electron Microscopy (TEM). After TEM observation, select high-purity components based on the TEM results (Figure 3B).

**Figure 3.**
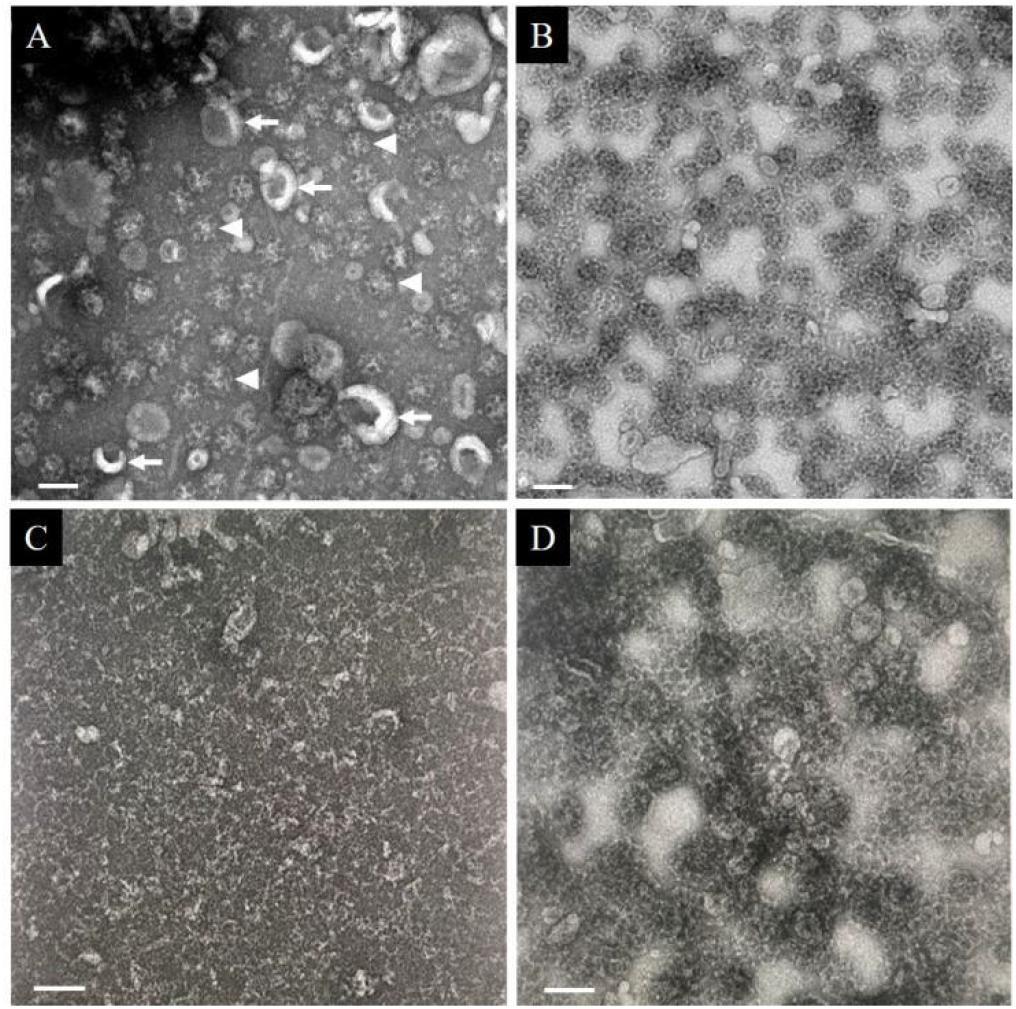
TEM analysis of samples during CCV purification and clathrin cage reconstruction. Representative TEM images of samples from sequential experimental steps. (A) Crude preparation prior to sucrose gradient centrifugation. Arrowheads mark CCVs and arrows mark exosomes. (B) Sample after continuous sucrose density gradient centrifugation. (C) Sample following clathrin depolymerization. (D) Purified, in vitro-reconstructed clathrin cages. Scale bar, 100 nm.

### 2.8 Depolymerization of clathrin cages and collection of SVs

Mix the selected samples with Buffer C at a 1:1 volume ratio and incubate at 4°C for 60 minutes. The mixture is then aliquoted into Beckman 11 x 34 mm round-bottom thick-walled ultracentrifuge tubes and centrifuged at 135,000 g for 20 minutes at 4°C using a Beckman Optima-Max-XP ultracentrifuge with an MLA-130 rotor. The pellet, which contains the SVs, is resuspended in an equal volume of Buffer A as used prior to depolymerization. The supernatant is dialyzed against Buffer D at 4°C overnight.

### 2.9 Reconstruction of the clathrin cage

The concentration was estimated by Sodium dodecyl sulfate polyacrylamide gel electrophoresis (SDS-PAGE). Concentrate the sample to at least 1 mg/ml using a 30 kDa MWCO ultrafiltration tube (Note 10). The concentrated sample was then stained with uranyl acetate for TEM to confirm that the clathrin cage had been completely depolymerized (Figure 3C).

Centrifuge the concentrated sample at 18,700 g for 10 minutes at 4°C and collect the supernatant (Note 11). Dialyze the supernatant against Buffer E at 4°C overnight. Next, centrifuge the dialyzed sample at 91,000 g at 4°C using a Beckman Optima-Max-XP ultracentrifuge with an MLA-130 rotor. Discard the supernatant and resuspend the pellet in 0.5 ml of Buffer E (Note 12) (Figure 2I). Centrifuge the resuspended sample at 18,700 g for 10 minutes at 4°C. Collect the supernatant and discard the pellet. Stain the supernatant with uranyl acetate, and confirm the reconstruction by TEM (Figure 3D).

### 2.10 Interactions between CCV, Auxilin and Hsc70

CCV, Auxilin + Hsc70, Hsc70 and CCV + Auxilin + Hsc70 were dialyzed overnight in Buffer A. The dialyzed samples were centrifuged at 134,000 g for 20 minutes at 4°C using the MLA-130 rotor. Each group was then resuspended in an equal volume of Buffer A. Western Blot (WB) was performed to determine the formation of the CCV-Auxilin-Hsc70 complex. ATP was added to CCV-Auxilin-Hsc70 complex and CCV alone to a final concentration of 1 mM, followed by incubation at room temperature for 30 minutes. TEM confirmed that the CCVs could be depolymerized with the assistance of Auxilin and Hsc70.

### 2.11 Analysis of vATPase

vATPase is a proton pump located on SVs. It consists of two subunits, V0 and V1. To confirm the presence of vATPase, we detected the V1 subunit in CCVs and SVs by WB. The vATPase activity SVs was measured using an ATP assay kit (Beyotime, S0026B). Lower luciferase units indicate greater ATP consumption. Since the assay kit contains background ATP, no additional ATP was added during the assay. Luciferase units were collected at 5 minutes after the start by Centro XS^3^ LB 960 of berthold Technologies.

### 2.12 SDS-PAGE analysis and western blot

The samples were separated using a 12% SDS-PAGE gel (1.5 mm thick, with a 4% stacking gel and a 12% resolving gel) and stained with Coomassie Brilliant Blue (BBI, Shanghai, CHN). The samples were separated by a 12% SDS-PAGE gel (0.75 mm-thick, 4% stacking, and 12% resolving). The separated samples were transferred from the SDS-PAGE gel to a PVDF membrane by wet transfer. The membrane was blocked with 5% non-fat milk in TBST for 1 hour. The membrane was then washed with TBST. Primary antibodies (anti-Clathrin heavy chain (ABclonal, A4943, 1:1000), anti-Auxilin (Proteintech, 21941-1-AP, 1:1000), anti-Hsc70 (ABclonal, A0415, 1:1000), anti-V1B (ABclonal, A6976, 1:1000), or anti-V1E (ABclonal, A3756, 1:1000)) were incubated overnight at 4°C. After washing with TBST, the membrane was incubated with Anti-Rabbit lgG-HRP (absin, abs20040, 1:10,000) for 2 hours at 4°C. Detection was performed using iBright 1500 of Invitrogen for imaging.

### 2.13 Transmission Electron Microscopy (TEM)

First, rinse the glow-discharged, carbon-coated copper grid with 3 μl of solution buffer and blot the excess solution from the edge of the grid using absorbent paper. Then, add 3 μl of the sample to the grid and incubate for 3 minutes, followed by blotting the solution dry from the edge with absorbent paper. Repeat the process of adding the sample and blotting dry. Next, add 3 μl of a 2% uranyl acetate solution to the grid for 20 seconds, then blot dry with absorbent paper. Repeat the application of the 2% uranyl acetate solution and blot dry after 60 seconds. The samples were visualized using a FEI Tecnai 12 electron microscope equipped with a LaB6 filament, operated at 120 kV. Images were captured with an FEI Ceta digital camera.

### 2.14 Notes

1. Dilute 10X Buffer A to the working concentration just before use.
2. Prepare all solutions immediately before use to ensure freshness and accuracy.
3. Keep pig brains on ice at all times to maintain maximum freshness.
4. Use blunt-tipped tweezers to gently peel off the fascia in large pieces to significantly reduce the processing time.Removing the fascia from the surface of the pig brain facilitates complete fragmentation. Focus only on removing the prominent fascia and avoid spending excessive time on this step. Prioritize shortening the purification time to maintain brain freshness. This step can be performed simultaneously with step 3.
5. Blend the sample for 5 seconds, followed by a 10-second pause, to prevent overheating. Use pre-cooled Buffer A and working in a 4°C environment helps keep the sample cool.
6. If the supernatant volume is insufficient, collect an equal amount from the symmetrical position of the centrifuge rotor to maintain balance. Retake the supernatant after 5 minutes, ensuring the centrifuge is properly balanced.
7. OptiSeal tubes must be fully filled with samples. Since the mixture density is significantly higher than water, use a sodium chloride solution or another high-density solution for balancing.Add ∼90% of the tube volume with water first, then gradually adjust the density by adding small amounts of sodium chloride until the appropriate balance is achieved. This adjustment compensates for varying mixture densities in different experiments.
8. Due to the toughness and stickiness of the precipitate, white resuspension often adheres to the tube wall. Resuspend the pellet with 1 mL of 1X Buffer A, then rinse the tube with another 1 mL of 1X Buffer A to increase the extraction rate.
9. Based on previous experiments, exosome peaks are observed at 20%-22.5% density, while CCV peaks appear at 22.5%-24% density (Supplementary Figure 1).
10. The success of assembly is highly dependent on concentration. Higher concentrations yield better results.
11. Check for any visible precipitation. Significant precipitation could result from prolonged sample storage or an excess of exosomes. Reassess the clathrin density to ensure sufficient concentration for successful reconstruction.
12. The precipitate forms small, transparent disc-like structures that are difficult to resuspend. Resuspend carefully to ensure most reconstituted clathrin cages are returned to the solution. Failure to fully resuspend the precipitate will severely reduce the yield.

## 3. RESULTS AND DISCUSSION

### 3.1 Purification of endogenous CCVs and in vitro reconstitution of clathrin cages

The overall workflow of this method is summarized in Figure 4. Fresh porcine brains were used as starting material; connective fascia was removed, and tissues were fully homogenized. Sequential differential centrifugation was applied to remove large poorly soluble contaminants, followed by Ficoll-sucrose gradient centrifugation to enrich vesicle fractions. Additional centrifugation steps were performed to deplete cytoskeletal contaminants, although exosomes still co-purified with CCVs in these samples. Taking advantage of the difference in sedimentation coefficients between CCVs and exosomes, we separated CCVs from exosomes using continuous sucrose density gradient centrifugation. Notably, samples with moderate purity can still be used for clathrin cage reconstitution, since residual exosomal contaminants are removed during the subsequent depolymerization and reconstitution steps.

**Figure 4.**
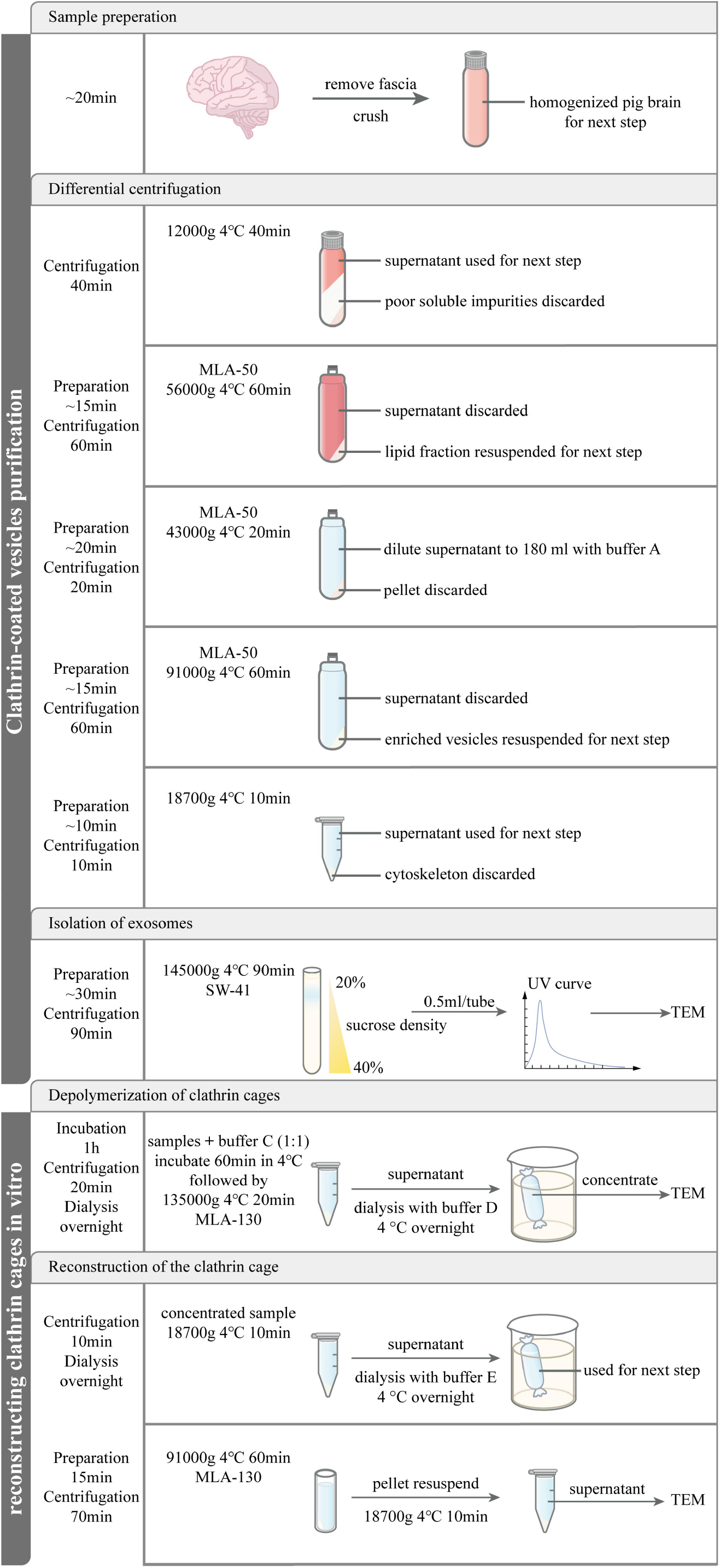
Schematic workflow for purification of endogenous CCVs and in vitro reconstruction of clathrin cages. Flowchart summarizing the full experimental procedure, including tissue homogenization, sequential differential and density-gradient centrifugation, clathrin cage depolymerization, dialysis, and subsequent in vitro reassembly of clathrin cages. Key centrifugation parameters, buffer exchange steps and downstream characterization (UV absorbance and TEM) are indicated.

Clathrin cages are sensitive to pH and ammonium ions. Therefore, purified CCV samples were dialyzed into alkaline Tris-containing buffer to trigger cage depolymerization. Smooth vesicles (SVs) released upon depolymerization exhibited low solubility and tended to aggregate. Lipid-containing particles were pelleted by ultracentrifugation, and the supernatant containing soluble clathrin and adaptor proteins (APs) was collected. After buffer exchange into assembly buffer, clathrin was reassembled into intact cages in the presence of APs.

Fractions collected across the purification workflow (S1, P1, S2, P2, S3, P3, S4, P4, S5 and P5) were analyzed by SDS-PAGE to monitor clathrin enrichment. The SDS-PAGE results showed a progressive increase in the relative abundance of the clathrin heavy chain (CHC) band, indicating gradual enrichment and purification of CCVs (Figure 5). Two commonly used Ficoll-sucrose concentrations (6.25% and 12.5% w/v) were compared in the vesicle enrichment step. No obvious difference in CCV yield or purity was observed from CHC band intensities on SDS-PAGE, suggesting both formulations are compatible with this protocol (Supplementary Figure 2).

**Figure 5.**
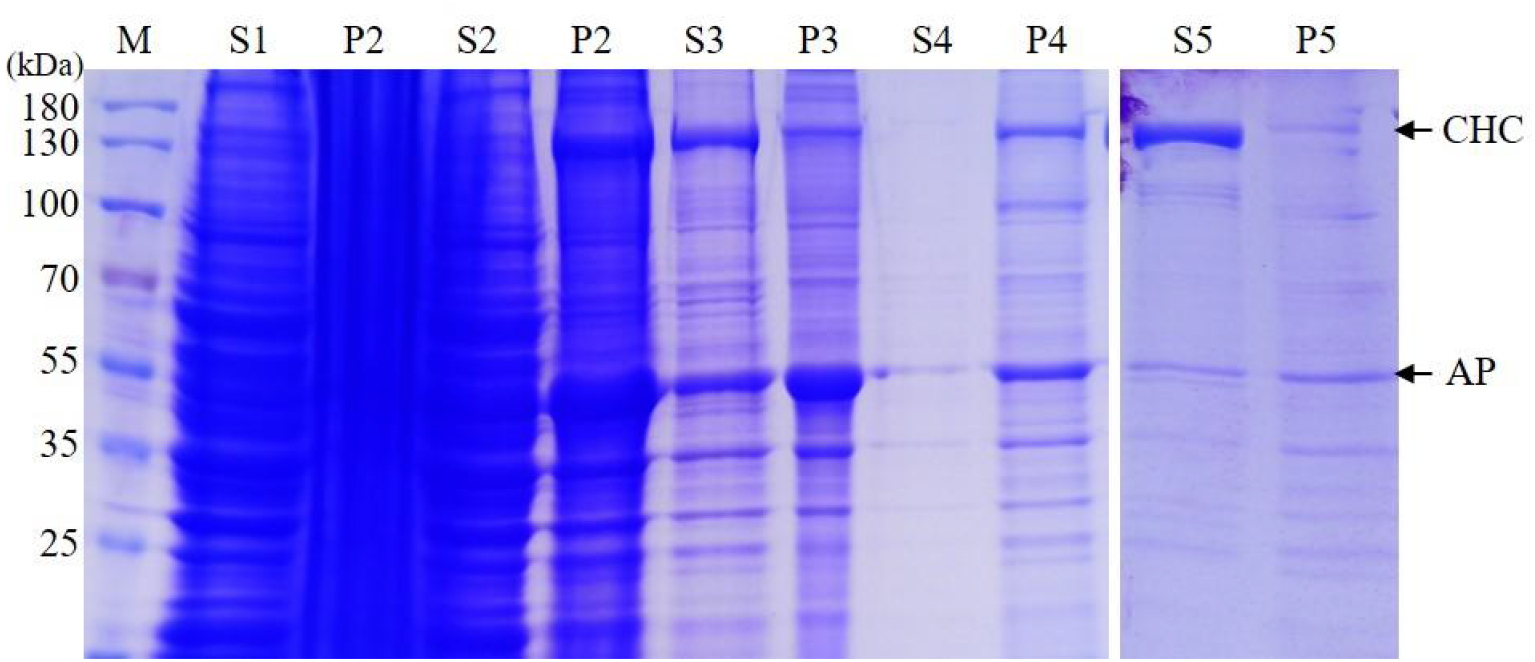
SDS-PAGE analysis of protein composition at successive purification stages. Coomassie-stained SDS-PAGE gel showing protein profiles of supernatant (S) and pellet (P) fractions collected across the purification workflow. Lane M: molecular weight marker (kDa). Arrows indicate clathrin heavy chain (CHC) and adaptor protein (AP).

Uranyl acetate negative-stain TEM of the S5 fraction confirmed substantial enrichment of endogenous CCVs, yet considerable exosomal contamination remained (Figure 3A). Consistent with their distinct sedimentation properties, continuous sucrose gradient centrifugation efficiently removed most exosomes from the CCV preparation (Figure 3B).

After buffer exchange into alkaline Tris buffer, TEM imaging confirmed complete depolymerization of clathrin cages after 60 min incubation at 4 °C (Figure 3C). Lipid contaminants including SVs and leftover exosomes were pelleted by ultracentrifugation, leaving soluble clathrin and APs in the supernatant. The protein mixture was concentrated and dialyzed against assembly Buffer E. Because reassembled clathrin cages have higher sedimentation coefficients than monomeric coat proteins, ultracentrifugation was used to harvest reconstituted cages. The pellet was resuspended and clarified by a short low-speed spin to remove large aggregates. TEM verified successful reconstitution of homogeneous clathrin cages free of encapsulated SVs (Figure 3D).

CCVs and exosomes differ in size and buoyant density: exosomes range from 30-150 nm with a density of 1.13-1.19 g/mL [14], while CCVs are 50-100 nm with a density of 1.20-1.25 g/mL [15,16]. These physical differences underpin the separation achieved by sucrose gradient centrifugation. Although this step was optimized for CCV purification, high-purity exosomes can also be recovered from the same gradient and may serve as material for separate exosome-focused studies.

For high-resolution structural biology, native endogenous CCVs are suboptimal. The encapsulated SVs interfere with cryo-EM reconstruction, and inherent heterogeneity of tissue-derived CCVs limits structural homogeneity. Our depolymerization-reconstitution pipeline circumvents these issues: all lipid-borne impurities (SVs and residual exosomes) are removed once the clathrin coat disassembles, allowing production of uniform, lipid-free clathrin cages suitable for structural analysis.

### 3.2 Functional validation of purified endogenous CCVs

Auxilin and Hsc70 are soluble proteins with low sedimentation coefficients and cannot be readily pelleted by ultracentrifugation alone. In contrast, intact CCVs sediment efficiently. When Auxilin and Hsc70 bind to CCVs, they co-sediment with the vesicles. Western blot results demonstrated stable complex formation between CCVs, Auxilin and Hsc70 (Figure 6A). Negative-stain TEM further confirmed ATP-dependent disassembly of clathrin coats upon incubation with Auxilin and Hsc70 (Figure 6B). Vacuolar-type ATPase (vATPase) is the proton pump resident on SV membranes. We confirmed the presence of the V1 subunit of vATPase by western blot (Figure 6C). ATP consumption assays verified retained enzymatic activity of vATPase on the SVs isolated from our CCV preparations (Figure 6D).

**Figure 6.**
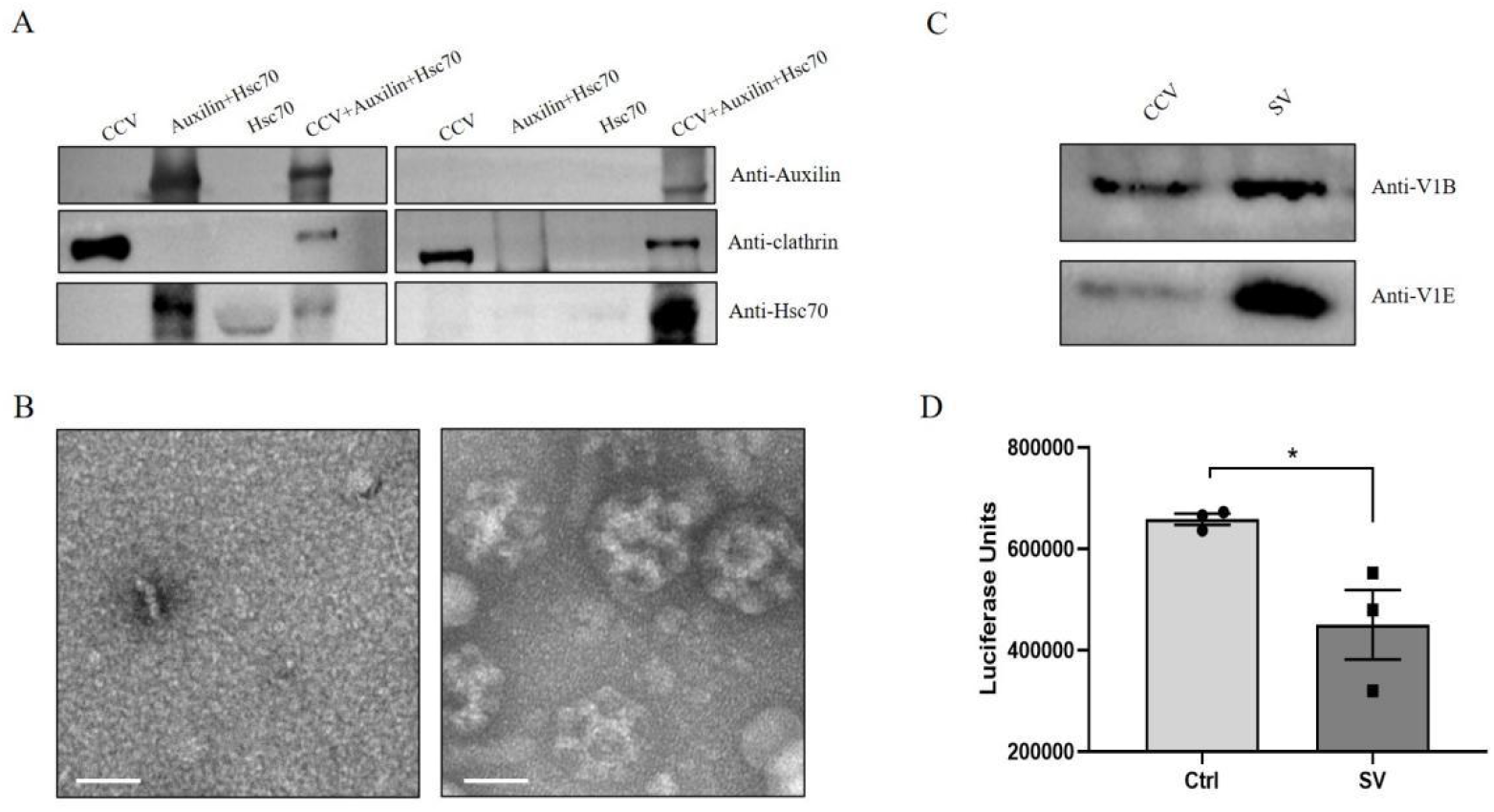
Functional and biochemical characterization of purified CCVs and smooth vesicles (SVs). (A) Western blots detecting auxilin, clathrin and Hsc70; Left: input proteins added into the reaction system; Right: proteins co-sedimented with clathrin-coated cages after ultracentrifugation. (B) Representative TEM images of CCV-auxilin-Hsc70 complex after 30 min ATP incubation (left) and CCVs alone (right). Scale bar, 50 nm. (C) Immunoblotting of vATPase V1B and V1E subunits in CCV and SV samples. (D) Luciferase signal from ATP assay after 5 min incubation with equal volumes of Buffer A or SV suspension. * denotes significant difference.

We also tested the stability of purified CCVs after cryopreservation. Samples were flash-frozen in liquid nitrogen, stored at -80 °C, and thawed after seven days. Visible fragmentation of clathrin cages was observed after thawing (Supplementary Figure 4). This indicates that the current preparation is not compatible with long-term frozen storage for structural studies, and fresh preparations are recommended.

Brain tissue was chosen as starting material due to its high abundance of CCVs, though it contains a complex mixture of other vesicle populations that necessitate multi-step purification. Tissues such as liver and kidney also contain CCVs. When using these alternative sources, the sucrose gradient separation step targeting exosomes may be simplified or omitted, shortening the overall purification workflow.

## 4. CONCLUSIONS

We describe a robust workflow for isolating functionally intact endogenous CCVs from porcine brain and reconstituting homogeneous, SV-free clathrin cages in vitro. The method separates CCVs from co-purified exosomes by exploiting differences in buoyant density and sedimentation behavior. A key advantage of this strategy is the depolymerization step, which removes intraluminal SV contaminants that cannot be eliminated by conventional gradient centrifugation alone. The purified CCVs retain physiological activity, including Auxilin-Hsc70-dependent uncoating and vATPase proton-pump activity on packaged SVs. The reconstituted clathrin cages are homogeneous and compatible with high-resolution cryo-EM and proteomic investigations. This protocol provides a useful tool for dissecting clathrin coat assembly, disassembly and clathrin-dependent membrane trafficking.

## Author contributions

Y.W., X.L., J.L., C.Y. and S.S. performed the experiments. Y.W., X.L., W.L., S.S. analyzed the data and wrote the manuscript. All authors contributed to the discussion of the data and editing the manuscript.

## Ethics approval and consent to participate

Not applicable.

## Data Availability Statement

The original contributions presented in this study are included in the article. Further inquiries may be directed to the corresponding authors.

## Declaration if Competing Interest

The authors declare that they have no competing interests.

