## Supplemental File for "Isolation of functional endogenous clathrin-coated vesicles and in vitro reconstitution of pure clathrin cages"

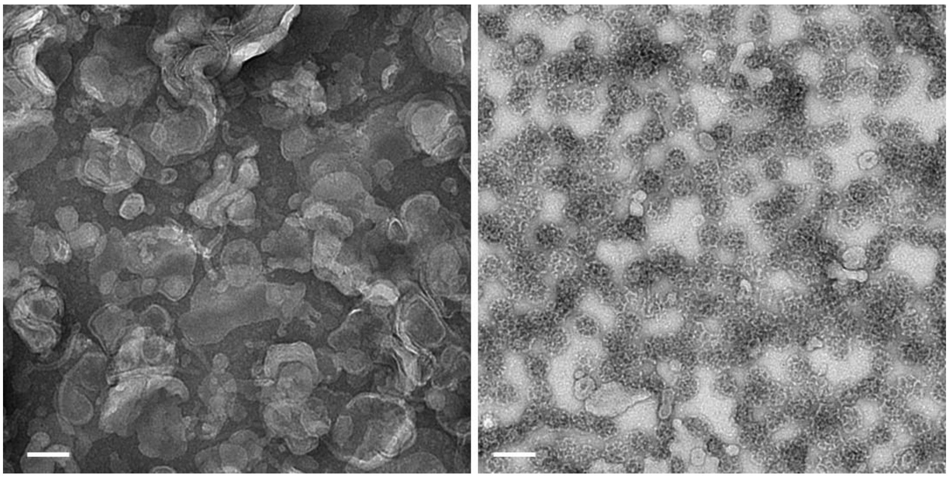

**Supplementary Figure 1. TEM micrographs for fractions collected from two sucrose gradient peaks.** Left: sample from the 20%–22.5% sucrose fraction; Right: sample from the 22.5%–24% sucrose fraction. Scale bar, 100 nm.

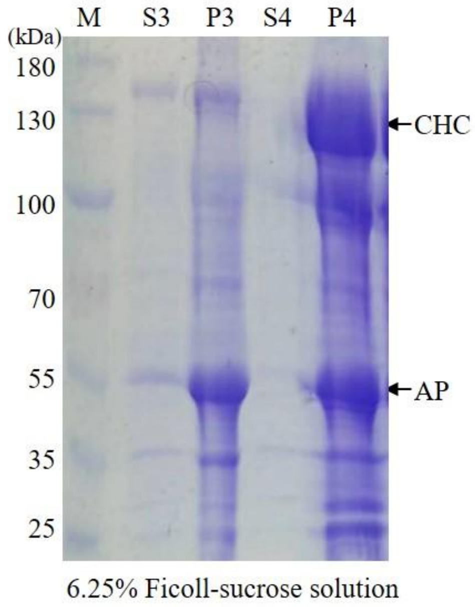


**Supplementary Figure 2. SDS-PAGE analysis of vesicle enrichment using 6.25% Ficoll-sucrose solution.** Coomassie-stained SDS-PAGE gel showing supernatant (S) and pellet (P) fractions. Lane M: molecular weight marker (kDa). Arrows indicate clathrin heavy chain (CHC) and adaptor protein (AP).

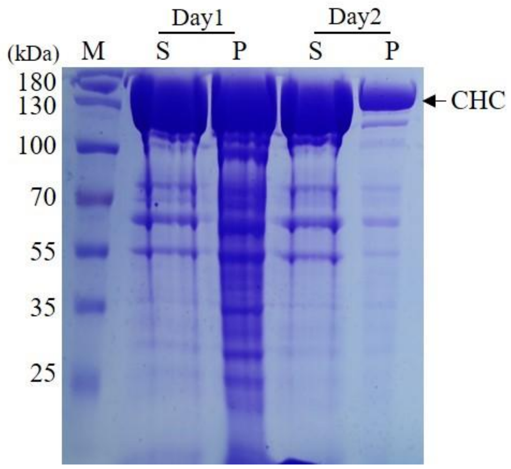


**Supplementary Figure3. SDS-PAGE analysis of clathrin samples without exosome depletion.** Coomassie-stained SDS-PAGE gel showing supernatant (S) and pellet (P) fractions of clathrin samples harvested on Day 1 and Day 2, when exosomes were not removed during purification. Lane M: molecular weight marker (kDa). Arrow indicates clathrin heavy chain (CHC).

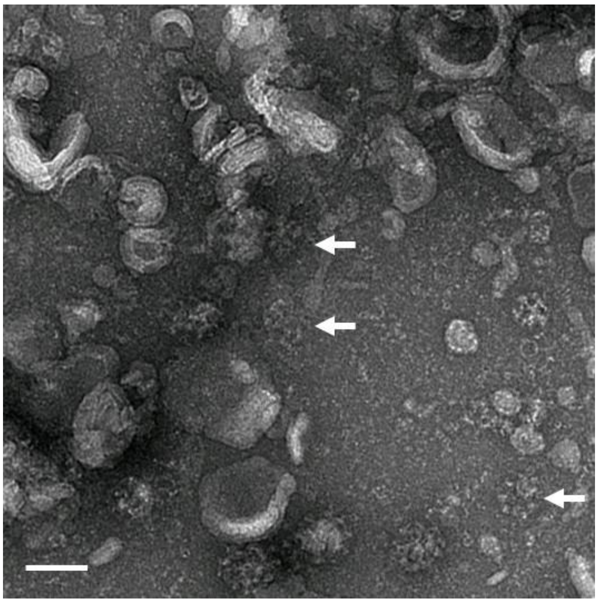

**Supplementary Figure 4. TEM characterization of clathrin cage samples after storage at −80 °C for seven days.** Representative TEM micrograph showing structural deterioration upon frozen storage. Arrows indicate broken clathrin cages. Scale bar, 100 nm.
